# All Models are Wrong, Some are Annotated: Automating Metadata in Biomedical Repositories

**DOI:** 10.64898/2026.04.23.720371

**Authors:** Inessa Cohen, Hongyi Yu, Robert A. McDougal

## Abstract

**Objective:** High-quality metadata is essential for scientific discovery, yet sparse annotations in rapidly growing repositories leave many biologically relevant details uncaptured. We evaluated whether large language models (LLMs) can accurately infer ion channel and receptor subtype metadata from source code in a neuroscience repository.

**Materials and Methods:** We extracted 5,133 model files from ModelDB. A subset of 1,100 was manually annotated; 253 were held out for testing, and the remainder split into training (80%) and validation (20%) sets. LLM-based approaches (GPT-5.2 and GPT-mini) were evaluated under zero-shot and heuristic-augmented prompting. Performance was assessed at type and subtype levels using accuracy, precision, recall, and F1 score. A feature-engineered XGBoost model using text- and simulation-derived features served as a baseline.

**Results:** LLMs outperformed the XGBoost baseline. At the type level, GPT-mini with heuristic augmentation achieved the highest performance (accuracy 96.0%, F1 0.962). At the subtype level, both GPT-5.2+heuristics and GPT-mini+heuristics achieved identical accuracy (88.1%), with GPT-5.2+heuristics achieving the highest F1(0.878). Model outputs were consistent across runs and errors confined to related mechanistic families.

**Discussion and Conclusion:** LLMs demonstrate strong potential for metadata annotation directly from source code, outperforming feature-engineering approaches with minimal tuning. However, performance varied across subtypes, and errors often reflected ambiguity or bias toward more common labels. These findings suggest LLMs may serve as practical tools for scalable metadata generation in biomedical repositories, although careful evaluation and domain-specific validation remain important. While demonstrated in computational neuroscience, this approach may generalize to repository-agnostic metadata annotation in other scientific code repositories.

## INTRODUCTION

Metadata is a cornerstone of modern scientific databases.^1^ By providing structured descriptions that capture what data represent, how they were generated, and where they were used, metadata enables raw files to be findable, interpretable, and standardized.^2^ Its importance is widely recognized in the FAIR data principles (Findable, Accessible, Interoperable, Reusable), which emphasize that machine-actionable metadata are essential to maximize scientific value and reproducibility.^3,4^ Metadata is especially critical in fields where models and datasets span multiple scales, including molecules, cells, circuits, behavior, as well as various formats such as code, images, and tables.^5^ It can document analytical approaches, code dependencies, and the biological context that defines a model’s scope and function.^6^ Without standardized metadata, valuable content may remain hidden from researchers harming reproducibility.^6^ These challenges are further amplified in the era of large language models (LLMs), where non-deterministic outputs and evolving computational workflows increase the need for transparent, well-structured, and reusable code and metadata.^7^

Despite the FAIR principles and recent NIH polices and guidance on metadata,^8^ the quality of metadata in online knowledge bases remains inconsistent. Manual curation is slow, prone to errors, and unable to keep pace with the rapid growth of open scientific repositories.^6^ In computational modeling, many repositories lack biologically meaningful annotations linking model components or parameters to their underlying code, making it difficult to retrieve, compare, and reuse models across studies.^9^ Thus, developing scalable approaches to infer biologically relevant metadata directly from model source code represents a significant step toward more intelligent and interoperable neuroscience databases.

Recent advances in large language models (LLMs) offer a new opportunity to automate metadata generation by leveraging pretrained knowledge and semantic understanding to identify biological and neural mechanisms.^10,11^ Since neuroscience mechanistic information is distributed across thousands of publications and code repositories, LLMs may be well suited to infer structural relationships implicitly described in language and code.^12^ Prior studies have demonstrated the potential of GPT-based models to automate curation and metadata extraction from the literature in computational neuroscience.^12^ Related work in molecular biology has also shown that language models can interpret high-dimensional biological data, such as transcriptomic profiles or T-cell receptor (TCR) sequences, to improve annotation and capture functional context.^10,11^

However, LLMs also have important limitations. They can generate factual errors or hallucinated associations, particularly when outputs are not grounded in empirical or mechanistic evidence.^13^ Prior work has shown that LLMs may exhibit inflexible reasoning, particularly in out-of-distribution scenarios, leading to errors in reasoning, instruction-following, and maintaining faithfulness to the provided context.^14,15^ These concerns are especially relevant for identifying biological mechanisms from source code, which may require domain-specific knowledge beyond simple pattern matching. Prompting strategies such as chain-of-thought and domain-specific heuristics may improve performance on multi-step inference tasks,^16^ but recent work suggests that improved final accuracy does not necessarily reflect valid intermediate reasoning that causally supports the conclusion.^17,18^ Thus, whether heuristic prompting meaningfully improves mechanistic reasoning in a biological inference remains uncertain.

Motivated by both the promise and limitations of LLMs, we investigated their ability to automate biologically meaningful metadata annotation in ModelDB (https://modeldb.science/), a large public computational neuroscience repository hosting source code from published neuroscience simulation studies.^19,20^ Many of these models implement ion channels and receptors using equations inspired by the classic Hodgkin and Huxley squid giant axon model.^21^ ModelDB’s metadata are often incomplete or imprecise because annotations are provided by model contributors at submission and are recorded at the study level rather than the file level, limiting their usefulness for defining the ground truth at the mechanism level. Automatically inferring biologically meaningful labels such as ion channel and receptor subtypes would enrich repository metadata with biological context and enhance the search, comparison, and reproduction of computational models.

Here, we evaluate whether modern LLMs can serve as robust classifiers for ion channel and receptor subtype annotation directly from source code. We compare GPT-based models against a feature-engineered XGBoost baseline, assess the impact of prompt heuristics and model size, and characterize error models across approaches. Our objective is to determine whether LLMs can provide accurate, scalable, and minimally tuned metadata annotation for mechanistic models, supporting practical deployment in biomedical repositories.

## MATERIALS AND METHODS

### Data Acquisition

A total of 14,730 NEURON mechanism files^22^ (.mod, written in NMODL) were downloaded from ModelDB in February 2025 (**Figure 1A**). After removing duplicates arising from comments and whitespace (via SHA-256 hash comparison), 5,133 unique model files remained. From these, 1,100 were randomly selected for manual annotation to generate the gold-standard outcome labels used for model development. All remaining unannotated files were retained for large-scale LLM inference and agreement analysis.

**Figure 1.**
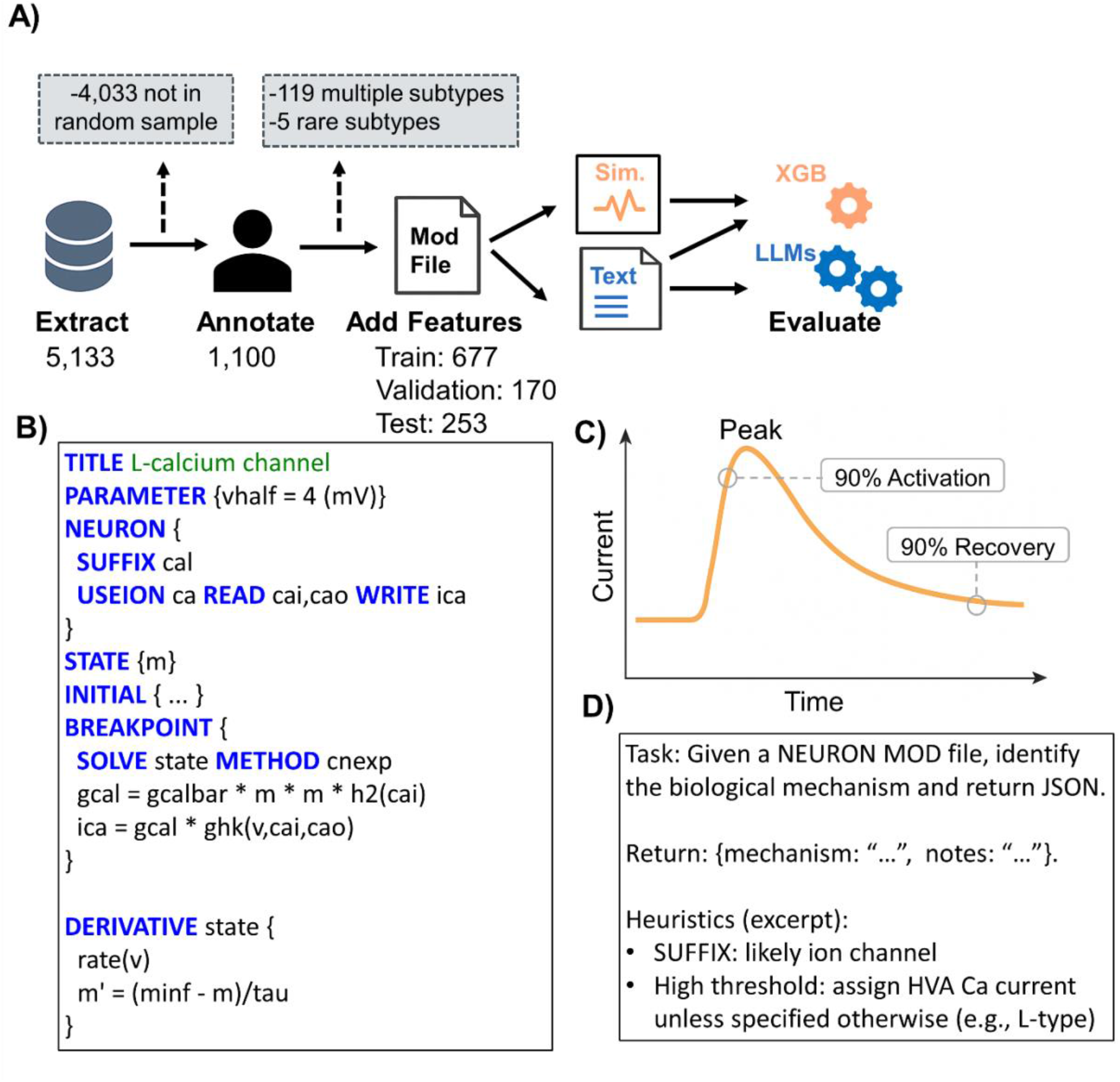
Methods Pipeline. (A) Workflow for extracting, annotating, and generating features from NEURON ModelDB model files, followed by XGBoost and GPT classification. Sample sizes are shown. Boxes show exclusion criteria. (B) Representative model file snippet of an L-type calcium channel with NEURON-specific syntax in capital letters. (C) Simulation traces provided dynamic features such as time to 90% activation and time to 90% recovery. (D) Excerpt of the heuristic-augmented prompt. Abbreviations: Sim., Simulated

### Outcome Label Annotation

The 1,100 files were manually annotated following ModelDB’s controlled vocabulary for ion channel and receptor mechanisms (https://modeldb.science/FindByCurrent). Each file was assigned to one of 19 ion channel and receptor subtypes, corresponding to the most granular subtype supported by its implementation (e.g., A-type potassium, GABA-A). Mechanisms that clearly did not represent any channel or receptor subtype (e.g., voltage clamps, artificial cells, gap junctions, calcium-accumulation mechanisms, utility files) were labeled as ‘Neither’. When subtype assignment was ambiguous, we examined how the mechanism interacted with membrane voltage and ionic variables in code and selected the closest matching subtype, favoring the most specific label available (e.g., a BK calcium channel was annotated as ‘BK’ rather than the broader category ‘calcium-activated potassium channel’). Confidence levels (low, medium, high) were recorded for each subtype assignment. Detailed annotation heuristics are provided in **Supplementary Table S1**. These annotations served as the reference standard for all model evaluations.

### GPT-based Label Prediction

Large language models (LLMs) were evaluated as the primary classification approach. Multiple GPT-based models were tested, including a full-capacity model (GPT-5.2) and a lightweight model (GPT-mini) to assess performance. Each NMODL file was provided as plain text with the 19-subtype vocabulary. Two prompting strategies were used: (1) a zero-shot baseline including only the file text and label list, and (2) a heuristics-augmented: prompt including the label list and concise domain-specific annotation guidelines derived from manual curation (e.g., NET_RECEIVE blocks, gating variable structure, etc., **Figure 1D, Supplemental Table S1, Appendix S1**). For each file, the full source text was submitted via API, and models were instructed to return a JSON object containing a mechanism label selected verbatim from the 19-category vocabulary and brief free-text notes. Responses were parsed and annotations were stored in a SQLite database. Files with missing JSON were flagged for reprocessing.

Performance was evaluated at both the subtype and type levels. Structured error analysis used human annotation as the reference standard and compared GPT-based models with the XGBoost model on the test set. Multiclass precision, recall, and F1 were computed using weighted averaging. To assess robustness and internal consistency, each GPT configuration was run multiple times, and pairwise agreement (Cohen’s kappa) was calculated across repeated runs, prompting strategies, and model sizes on the full corpus of 5,133 mod files. To further characterize model disagreement, we examined the annotated subset of 1,100 mod files for which human confidence ratings were available.

### XGBoost Baseline Model

As a mechanistic baseline comparator, we developed a multiclass XGBoost classifier using text-based and simulation-derived features (**Figure 1A-C**). Text-based features were primarily binary indicators capturing NMODL-specific syntax (e.g., SUFFIX, POINT_PROCESS, NET_RECEIVE, ion usage, gating variables, **Figure 1B**). Simulation-derived features were obtained by instantiating each mechanism in a single-compartment NEURON model under a standardized three-step voltage protocol: a -65 mV hold, 1,000 ms to a test voltage, and clamp release to observe the channel response. From these traces, summary statistics (e.g., extrema, initial/final values, and time-to-90% activation and recovery) were extracted to represent mechanistic behavior (**Figure 1C**).

A subset of 253 files was reserved for testing (**Figure 1A**). The remaining files were stratified by subtype into training (80%) and validation (20%) sets. A single subtype-level XGBoost model was trained to predict 19 subtypes, and higher-level type labels were derived from the predicted subtype labels. Features were preprocessed in Python 3.10 using the Feature-engine package, including feature selection, imputation, normalization, and correlation pruning, and hyperparameters were optimized via randomized cross-validation (**Supplemental Table S2**). Files were excluded if they had multiple subtypes, could not be processed reliably (e.g., pointer files or unavailable files), or had low-confidence annotations. Rare subtype labels with counts below 10 were consolidated where possible and excluded if counts remained below 10 after collapsing. Performance was evaluated on the external test set using confusion matrices, accuracy, precision, recall, and F1 score metrics.

## Data/Code Availability

The code used to generate the results and analyses in the study is publicly available on GitHub at: https://github.com/innacohen/mod-annotation.

## RESULTS

### Dataset Characteristics

Of the 1,100 annotated files, 693 (63%) were ion channels, 134 (12.2%) were receptors, and 273 (24.8%) were neither. Subtype distributions were similar across training, validation, and test sets (**Table 1**). Potassium channels were the most common type (30.3%), especially A-type, delayed rectifier, and calcium-activated subtypes.

**Table 1.** Dataset characteristics. Counts and percentages of ion channel and receptor mechanism categories in the training, validation, and test sets.

|  | **Train**<br>**(N=677)** | **Validation**<br>**(N=170)** | **Test**<br>**(N=253)** |
| --- | --- | --- | --- |
| **Type** |  |  |  |
| I Ca | 71 (10.5%) | 17 (10.0%) | 36 (14.2%) |
| I H | 31 (4.6%) | 8 (4.7%) | 8 (3.2%) |
| I K | 207 (30.6%) | 53 (31.2%) | 74 (29.2%) |
| I Na | 93 (13.7%) | 23 (13.5%) | 36 (14.2%) |
| I Other | 22 (3.2%) | 6 (3.5%) | 8 (3.2%) |
| Neither | 166 (24.5%) | 42 (24.7%) | 65 (25.7%) |
| Receptor | 87 (12.9%) | 21(12.4%) | 26 (10.3%) |
| **Subtype** |  |  |  |
| I Ca (HVA) | 45 (6.6%) | 11 (6.5%) | 22 (8.7%) |
| I Ca (Rare) | 9 (1.3%) | 2 (1.2%) | 3 (1.2%) |
| I Ca (T-type LT) | 17 (2.5%) | 4 (2.4%) | 11 (4.3%) |
| I H | 31 (4.6%) | 8 (4.7%) | 8 (3.2%) |
| I K (A-type) | 61 (9.0%) | 15 (8.8%) | 19 (7.5%) |
| I K (Ca-activated) | 45 (6.6%) | 12 (7.1%) | 12 (4.7%) |
| I K (Delayed Rectifier) | 48 (7.1%) | 12 (7.1%) | 22 (8.7%) |
| I K (M-type) | 24 (3.5%) | 6 (3.5%) | 10 (4.0%) |
| I K (Rare) | 29 (4.3%) | 8 (4.7%) | 11 (4.3%) |
| I Na (Persistent) | 13 (1.9%) | 3 (1.8%) | 6 (2.4%) |
| I Na (Rare) | 13 (1.9%) | 3 (1.8%) | 8 (3.2%) |
| I Na (Slow inactivation) | 13 (1.9%) | 3 (1.8%) | 3 (1.2%) |
| I Na (Transient) | 54 (8.0%) | 14 (8.2%) | 19 (7.5%) |
| I Other (Rare) | 22 (3.2%) | 6 (3.5%) | 8 (3.2%) |
| R GABA | 23 (3.4%) | 6 (3.5%) | 8 (3.2%) |
| R Glutamate (AMPA) | 25 (3.7%) | 6 (3.5%) | 5 (2.0%) |
| R Glutamate (NMDA) | 26 (3.8%) | 6 (3.5%) | 10 (4.0%) |
| R Other (General) | 13 (1.9%) | 3 (1.8%) | 3 (1.2%) |
| Z Neither | 166 (24.5%) | 42 (24.7%) | 65 (25.7%) |

### LLM Performance

GPT-based models outperformed the XGBoost baseline at type-level classification (**Table 2**). Accuracy was identical for GPT-5.2 with and without heuristic augmentation (0.953), although heuristics slightly reduced precision (0.962 to 0.960) and F1 (0.954 to 0.953). GPT-mini achieved performance comparable to the full-size model (accuracy = 0.945, F1=0.947). With heuristic augmentation, GPT-mini outperformed all other models, achieving the highest overall accuracy (0.960) and F1 score (0.962).

**Table 2.** Type-Level Performance Metrics. Performance metrics for type-level mechanism classification across models, including accuracy and weighted metrics (precision, recall, and F1 score). Bolded row shows highest performing model.

| Model | Accuracy | Precision | Recall | F1 |
| --- | --- | --- | --- | --- |
| GPT-5.2 | 0.953 | 0.962 | 0.953 | 0.954 |
| GPT-5.2 + heuristics | 0.953 | 0.960 | 0.953 | 0.953 |
| GPT-mini | 0.945 | 0.956 | 0.945 | 0.947 |
| **GPT-mini + heuristics** | **0.960** | **0.969** | **0.960** | **0.962** |
| XGB | 0.909 | 0.914 | 0.909 | 0.911 |

At the subtype-level, GPT-based models again outperformed XGBoost (**Table 3**). GPT-5.2 achieved an accuracy of 0.862 (F1=0.862), which improved with heuristic augmentation (accuracy=0.881, F1=0.878). GPT-mini showed slightly lower baseline performance (accuracy=0.858, F1=0.846), and improved with heuristic augmentation (accuracy=0.881, F1=0.862), although it did not surpass GPT-5.2 with heuristics. A small number of subtypes resulted in failed predictions, ranging from 1 to 3 across models (**Table 3**). Inspection of these subtypes indicated that they were primarily associated with ambiguous classes including Neither, R Other (General), and I Other (Rare).

**Table 3.**
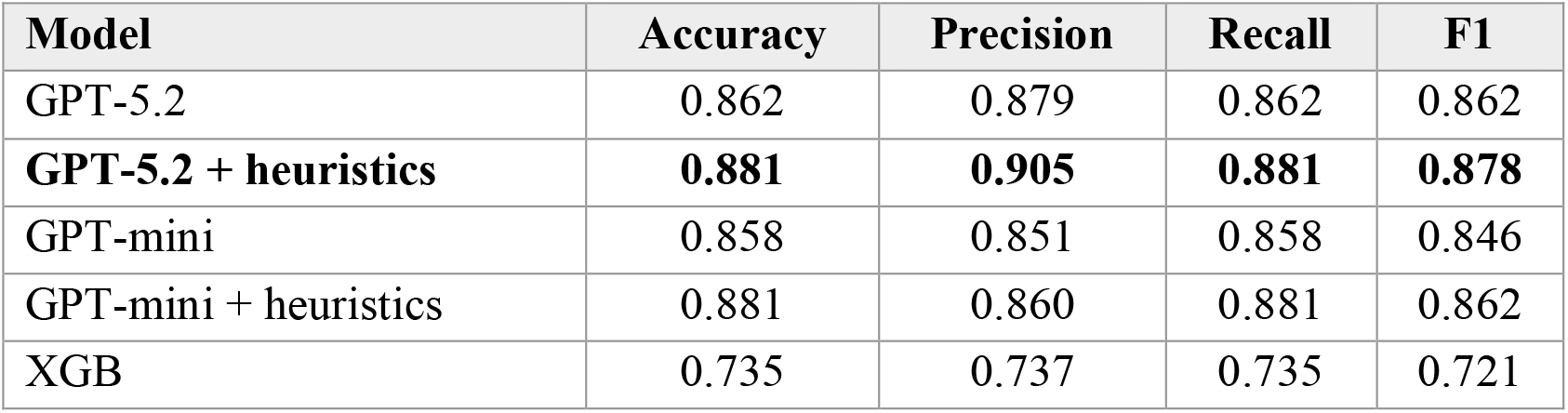
Subtype-Level Performance Metrics. Performance metrics for subtype-level mechanism classification across models, including accuracy and weighted metrics (precision, recall, and F1 score). Bolded row shows highest performing model.

Subtype-level patterns showed that heuristic augmentation produced small shifts in true positive rates for a limited number of subtypes, while most mechanisms exhibited tied performance between baseline and heuristic models (**Figure 2A-D**). For both GPT-5.2 and GPT-mini, sensitivity improved for selected Na and K channel subtypes, although these changes were modest (**Figure 2A, 2C**). False positive rates varied more substantially across subtypes, with the highest variability observed in rare and ambiguous classes (**Figure 2B, 2D**). XGBoost achieved perfect sensitivity for Ca (HVA) and I Na (Slow inactivation), while GPT-based models achieved perfect sensitivity for twelve subtypes, particularly among subtypes with small sample sizes. Specificity was consistently high across all models (range: 95.7-100%) with minimal variation across subtypes.

**Figure 2.**
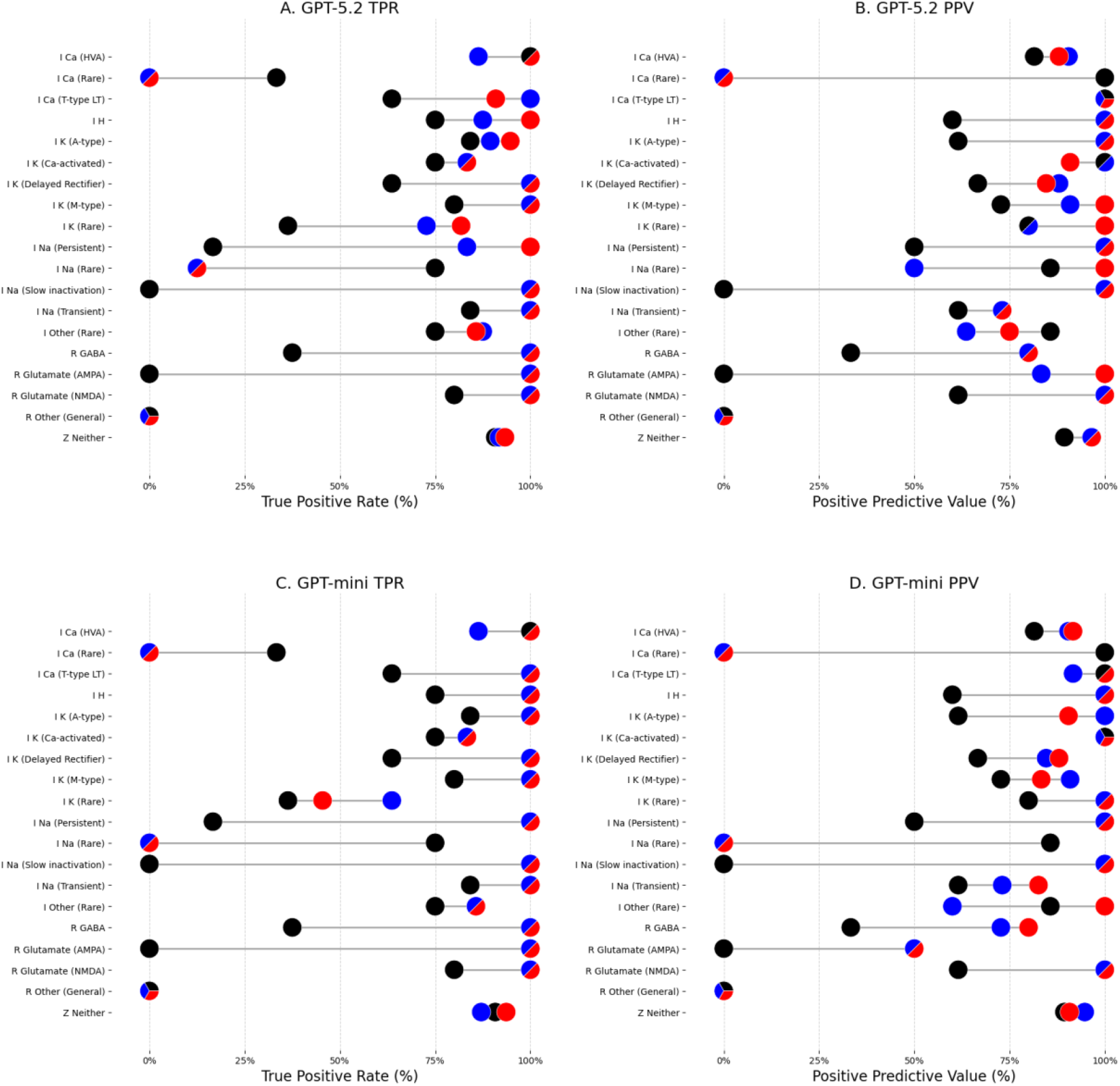
Subtype-level performance. Top row shows true positive rate (A) and positive predictive value (B) for GPT 5.2; bottom row shows corresponding results for GPT-mini (C-D). Each point represents performance for an individual subtype; ties between models are shown as multi-colored points. Colors of true instances per subtype are shown on the right. Colors: XGB (black), GPT (blue), GPT with heuristics (red).

### Agreement Across LLM Runs and Models

Agreement between repeated runs of the same full-size model (GPT-5.2) was high, with approximately 4.6% of files receiving discordant labels across repeated runs (kappa=0.95). Agreement decreased modestly when comparing different prompting strategies within the same model (8-9% disagreement), and more substantially when comparing different model variants (approximately 11.4-13.6% disagreement across GPT-5.2 and GPT-mini, **Supplemental Table S3**). This agreement was not meaningfully associated with human annotation confidence, with similar agreement rates observed for high-confidence vs. lower-confidence annotations (95.3% vs. 94.4%, **Supplemental Table S4**). In contrast, agreement decreased when comparing different model sizes and prompting strategies. For example, agreements between GPT-5.2 and GPT-mini were substantially higher for high-confidence annotations (90.3% vs. 79.7%). A similar pattern was observed when comparing GPT-mini with and without heuristic prompting (91.0% vs. 81.9%).

### LLM Error Analysis

Of the 253 files in the test set, 49 files were misclassified by at least one GPT model (**Figure 3**). Of those, 22 were misclassified across all GPT variants, whereas the remainder had model-specific errors. GPT-mini with heuristic augmentation showed more coherent, within-family misclassifications compared to XGBoost that demonstrated more cross-family errors, including greater spillover between ion channel and receptor classes (**Figure 4**).

**Figure 3.**
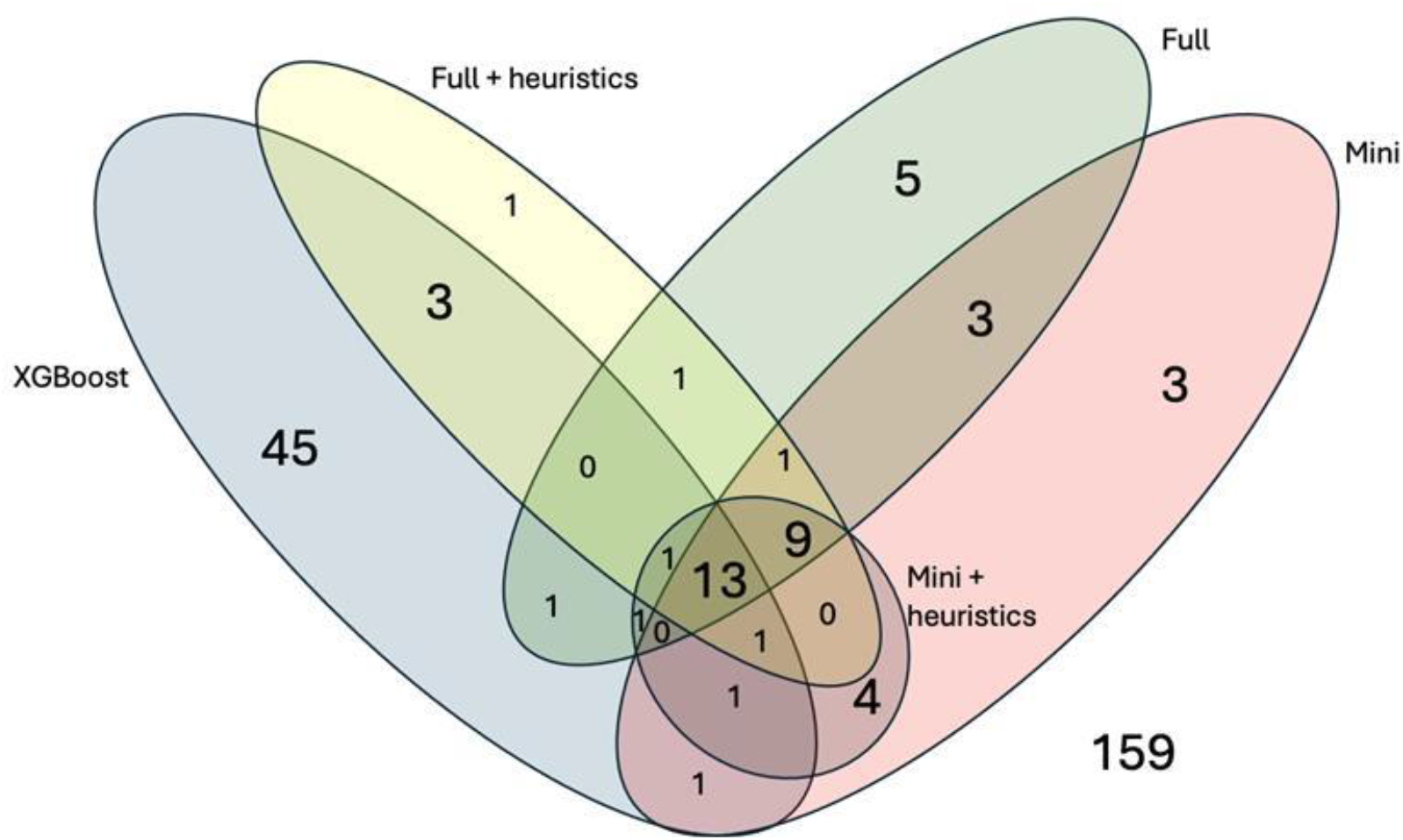
Error overlap across models. Venn diagram showing misclassified cases for XGBoost, GPT-mini, GPT-5.2, and their heuristic variants. Overlaps indicate shared errors; non-overlapping regions indicate model-specific errors. Outside (n=159) represents cases correctly classified by all models.

**Figure 4.**
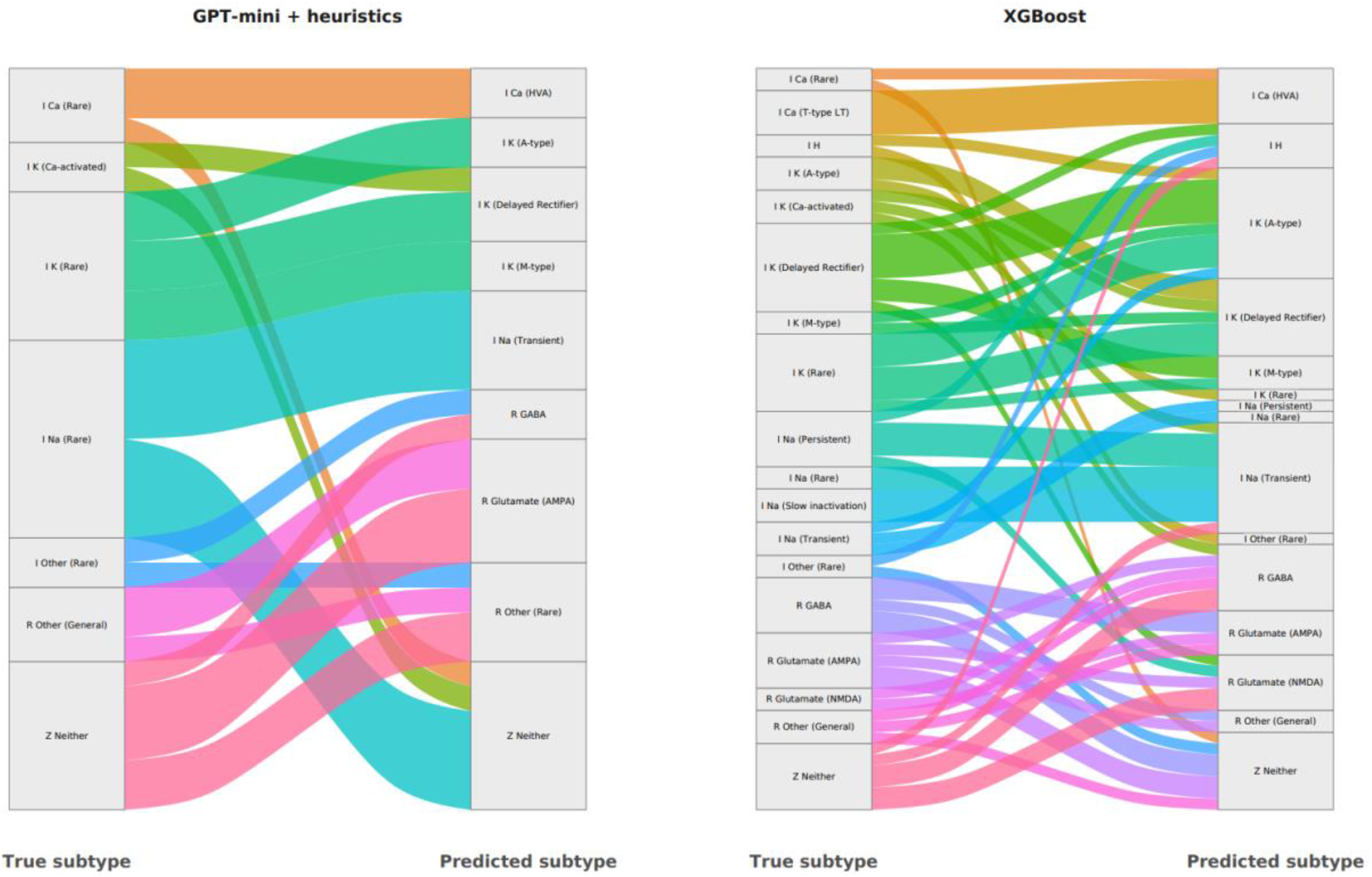
Subtype-level errors patterns. Sankey diagram depicting flows from true subtype to predicted subtype for GPT-mini with heuristic augmentation (left panel) and XGBoost (right panel). Only errors are shown.

Across all models, 159 of 253 (62.8%) subtypes were correctly classified by every model. In 45 cases (17.8%), at least one GPT model correctly classified a file that the XGBoost model misclassified. In contrast, XGBoost alone correctly classified 9 cases (3.6%) in which all GPT variants were incorrect (**Supplemental Figure S2**). Several LLM outputs included multiple predicted mechanisms for a single mod file, despite prompts requesting a single label.

Disagreements among GPT models, and between GPT and human annotation were clustered within closely related mechanistic families (e.g., I Na Transient vs. I Na Rare), and ambiguity between receptor-based and current-based mechanisms (e.g. I Other Rare vs. R Other Rare, **Figure 4**).

### Top Features (XGBoost Baseline)

As a baseline comparator, we examined feature importance for the XGBoost model (**Supplemental Figure 3**). Split-based importance emphasized simulation-derived dynamics, while SHAP distributed importance across both simulation and text-derived features. These results indicate that XGBoost primarily relied on voltage-dependent properties with supplemental signal from code structure, providing a mechanistic baseline against which LLM performance was evaluated.

## DISCUSSION

We found that GPT-based models consistently outperformed a feature-engineered XGBoost baseline for both type-level and subtype-level ion channel classification, while requiring minimal prompting and maintaining strong performance after model downsizing. Lightweight GPT-mini models achieved accuracy comparable to the full-capacity GPT, and heuristic augmentation produced only modest gains, primarily for less common subtypes. Together, these results suggest that modern LLMs can function as robust scientific classifiers in a largely zero-shot setting.

Although XGBoost relied primarily on simulation-derived dynamic features (e.g., activation, recovery, voltage-dependent kinetics), GPT models achieved superior performance directly from source code alone. This highlights a key distinction between approaches: feature-engineered models depend on explicitly constructed representations of mechanism behavior, whereas LLMs implicitly integrate contextual cues from variable names, comments, and code structure. The strong performance of GPT without access to simulation-derived features indicates that pretrained language models already encode substantial domain knowledge relevant to mechanistic classification. These findings align with recent work demonstrating that LLMs can perform biomedical classification and appraisal tasks in zero-shot settings with performance comparable to fine-tuned models, while providing interpretable justifications for their predictions.^23^

Zero-shot GPT-based models performed comparatively well on rare or poorly defined subtypes, such as potassium and receptor models lacking dynamic features, by leveraging contextual cues within the source code. When augmented with domain-specific guidelines, GPT-based models achieved the highest overall performance, with lightweight models performing best at the type level and full-capacity models achieving the highest performance at the subtype level, exceeding both XGBoost and GPT without heuristics. However, gains from heuristic augmentation were modest relative to the base model. This pattern suggests diminishing returns from additional prompting as model capacity increases, with the magnitude of improvement varying across models and tasks, and full-capacity GPT achieving the highest subtype-level performance. These findings suggest that additional rules primarily help resolve edge cases rather than substantially altering performance.

Beyond aggregate accuracy, agreement analyses provided additional insights into the reliability of LLM-based annotation. Repeated runs of GPT-5.2 showed near-perfect agreement, indicating high internal consistency and aligning with prior work showing that LLMs often converge on similar answers across multiple sampled reasoning paths.^18^ In contrast, agreement decreased when comparing different model sizes and prompting strategies, suggesting that cross-model variability reflects systematic differences in representation rather than random noise.

Differences between full-capacity and lightweight GPT models appeared greater in files with lower-confidence human annotations, suggesting that some model disagreement tracks underlying ambiguity in the data.^24^ Files with high-confidence human labels showed higher cross-model agreement, whereas lower-confidence cases were more likely to produce disagreement. This pattern suggests that at least some LLM disagreements may highlight genuinely ambiguous cases that warrant further review, rather than simply reflecting model failure.

Error analyses further revealed that disagreements among GPT models and between GPT and human annotation were not uniformly distributed across labels, but instead clustered within closely related mechanistic families, including closely related sodium subtypes and ambiguity between receptor-based and neither-type mechanisms. These patterns reflect ontological ambiguity inherent in labeling rather than arbitrary classification, indicating that LLM errors are structured and biologically adjacent.

Interpretation of misclassification warrants caution, as mechanism labels impose discrete categories on models that may exhibit biological properties. Some errors occurred between closely related subtypes, suggesting that predictions may reflect partial mechanistic correctness rather than complete misclassification. In addition, we observed a more systematic tendency for LLMs to favor more prevalent or canonical subtypes when classifying ambiguous mechanisms, rather than assigning rarer or less well-defined categories, suggesting a bias toward high-frequency labels under uncertainty. Preliminary ablation experiments, in which descriptive elements were removed from source code, further suggest that models can rely on structural features beyond textual cues, although these analyses were limited in scale. These findings highlight the need for evaluation frameworks that account for biological similarity rather than exact-match accuracy.

Taken together, these findings support the view that LLMs can serve as scalable, reliable annotators of mechanistic metadata, with performance that reflects underlying biological structure rather than arbitrary pattern matching. At the same time, the use of LLMs introduces new challenges for reproducibility and transparency, as model outputs may vary across runs and versions, reinforcing the need for well-structured, well-documented, and reproducible computational workflows.^7^

Our approach has several limitations. First, many model files are intrinsically ambiguous because mechanisms may mix channel-like and receptor-like behaviors, contain vestigial code, rely on naming conventions that do not reflect the implemented equations, or encode processes such as calcium pumps that generate current but do not fit cleanly into ion-channel categories, making the ground truth itself uncertain. As a result, files containing more than one mechanism of subtype were excluded, simplifying the task but limiting applicability to single mechanisms.

Second, model performance was affected by class imbalance: most files belonged to the Neither category, which is highly heterogenous. Although collapsed into a single class for modeling, these mechanisms remain meaningful, and this simplification likely obscured structure that a richer taxonomy could capture. In addition, sensitivity was 100% for several subtypes, although these estimates were derived from limited sample sizes and may be less precise. Finally, simulation-derived features were only available for mechanisms that generated ionic currents, limiting performance of receptor-like and Neither mechanism, and several rare subtypes (e.g. chloride channels) appeared too infrequently for supervised learning.

Future work includes expanding simulation-based features to receptors, accommodating mixed and rare subtypes, developing hierarchical label structures, evaluating generalizability to other simulators, and integrating the workflow into the ModelDB submission and browsing interface. In addition, alternative evaluation frameworks could be explored that move beyond exact-match accuracy by awarding partial credit for biologically similar predictions. For example, confusing delayed-rectifier and A-type potassium channels, both potassium currents with overlapping dynamics, may be less severe than misclassification across unrelated mechanism classes.

Distance-aware or hierarchy-aware scoring may yield more informative evaluations. In addition to improving single-model performance, a natural extension of this work is the use of agent-based frameworks, in which multiple specialized models evaluate complementary aspects of mechanistic code and a coordinating agent integrates their outputs. Such approaches may better capture ambiguity and improve robustness, particularly for mixed or uncertain subtype classifications.

Lightweight GPT models demonstrated performance comparable to full-capacity models at substantially lower computational cost, suggesting that GPT mini may be preferred for large-scale deployment. Full GPT models demonstrated near-identical performance between runs regardless of heuristics suggesting that LLMs can be used reliably out of the box with minimal prompting. A confidence-guided pipeline, where the classifier defers to GPT, when unsure, and vice-versa, may better reflect complementary strengths of mechanistic and language-based models and reduce misclassification on rare or ambiguous subtypes.

Overall, this work demonstrates that modern LLMs can provide accurate, robust, and minimally tuned metadata annotation directly from mechanistic source code. While LLMs introduce new challenges for reproducibility, by enabling scalable subtype labeling without extensive feature engineering or prompt design, LLM-based approaches offer a practical path toward metadata automation in biomedical repositories and may help align repository content more closely with FAIR data principles.^3,4^

## Supplementary File

**Appendix S1**. Full LLM Prompt Templates

### GPT 5.2 / GPT mini

You are an expert in computational neuroscience. The following is a MOD file from a NEURON computational neuroscience model. Briefly discuss the biology that the above code appears to be modeling. End by generating JSON of the form {‘mechanisms’: [‘term1’, ‘term2’], ‘type of model’: ‘free text’, ‘notes’: ‘any notes of unusual things in the model’} where the terms (typically 0 or 1 but could be more represent the mechanism(s) (currents, pumps, and receptors) that are modeled in the file, chosen verbatim from the list below. When interpreting this list, HVA is high voltage activated, LT is low threshold, Rare should be interpreted as any other type of that channel that isn’t covered by a more specific channel, ‘I Other (Rare)’ is a current (e.g., Cl) that is not a Ca, K, or Na current, a prefix of R indicates a receptor (e.g., ‘R Other (Rare)’ is a synaptic receptor that is not GABA, AMPA, or NMDA), and ‘Z Neither’ is something that is neither a receptor nor a current (this could be a pump or it could be a utility file with many VERBATIM blocks that has no direct biological interpretation). Provide no additional explanation after the JSON.’I Ca (HVA)’, ‘I Ca (Rare)’, ‘I Ca (T-type LT)’, ‘I H’, ‘I K (A-type)’, ‘I K (Ca-activated)’, ‘I K (Delayed Rectifier)’, ‘I K (M-type)’, ‘I K (Rare)’, ‘I Na (Persistent)’, ‘I Na (Rare)’, ‘I Na (Slow inactivation)’, ‘I Na (Transient)’, ‘I Other (Rare)’, ‘R GABA’, ‘R Glutamate (AMPA)’, ‘R Glutamate (NMDA)’, ‘R Other (Rare)’, ‘Z Neither’

### GPT 5.2 + heuristics / GPT mini + heuristics

You are an expert in computational neuroscience. The following is a MOD file from a NEURON computational neuroscience model. Briefly discuss the biology that the above code appears to be modeling. You can use the following heuristics if it helps you to understand:

NET_RECEIVE: Likely receptor; discrete events rather than continuous current.

POINT PROCESS: Likely receptor but could be some other localized mechanism (e.g. neither - graded synapse, neither - gap junction).

SUFFIX: Likely ion channel but could be neither (e.g., SUFFIX NOTHING).

WRITE: intracellular concentrations only Not an ion channel; usually ‘Neither’ (e.g., WRITE cai calcium accumulation mechanism)

Other terms that indicate neither type ARTIFICIAL CELL, Clamp, ELECTRODE, excessive installs/VERBATIM blocks, POINT_PROCESS gap, i=ic

File name / comments: Not reliable; ‘synaptic current’ in comments ≠ receptor model could be a graded synapse; commented out names like ‘AMPAA’ in a ‘GABAA’ model Fast K: Ambiguous; may mean K-A, K-DR, or K-UR → need context.

A-type: Default = A-type transient unless ‘slow’; may appear as Kv4, Afast, transient outward current, ITO.

K slow: Assign K-slow unless explicitly marked A-type slow or M-slow. HH variant: Usually NaT or K-DR (sometimes K-A).

TTX sensitive: Assign NaT.

Anomalous rectifier / I-Funny / I-H: Assign I-H, not K-IR.

High threshold / High voltage: Assign HVA Ca current unless otherwise specific subtype specified (e.g., L-type).

Ligand-dependent: Likely a receptor (e.g, state transitions depend on neurotransmitter concentration)

Post-synaptic voltage-dependent: Likely an ion channel (e.g., depends on gating variables m/h/n)

Presynaptic voltage-dependent: Likely R Other (Rare) – graded synapse (e.g., conductance depends on vpre without NET RECEIVE)

Sodium, 1 m-gate: NaP.

Sodium, 1 h-gate: Ambiguous (flag).

Sodium, 2 gates: NaT (unless otherwise labeled, e.g., NaV1.9 → NaP).

Sodium, 3 gates: Na with slow inactivation.

Potassium, 1 n-gate: K-DR (Kv2/Kv3), unless explicitly labeled Kv1.1 (low threshold). Potassium, 2 gates: K-A, unless explicitly Kv2.x → K-DR.

Potassium, 3+ gates: General K; refine if Kv4.x → A-type.

NaV1.9: NaP.

Kv1.x (e.g., 1.1, 1.2, 1.5, Shaker): I_KLT or D-type. Kv2: K-DR.

Kv3: High-voltage K-DR.

Kv4 / Shal: A-type / ITO.

Kv7.x / KCNQ: M-type.

End by generating JSON of the form {‘mechanisms’: [‘term1’, ‘term2’], ‘type of model’: ‘free text’, ‘notes’: ‘any notes of unusual things in the model’} where the terms (typically 0 or 1 but could be more represent the mechanism(s) (currents, pumps, and receptors) that are modeled in the file, chosen verbatim from the list below. When interpreting this list, HVA is high voltage activated, LT is low threshold, Rare should be interpreted as any other type of that channel that isn’t covered by a more specific channel, ‘I Other (Rare)’ is a current (e.g., Cl) that is not a Ca, K, or Na current, a prefix of R indicates a receptor (e.g., ‘R Other (Rare)’ is a synaptic receptor that is not GABA, AMPA, or NMDA), and ‘Z Neither’ is something that is neither a receptor nor a current (this could be a pump or it could be a utility file with many VERBATIM blocks that has no direct biological interpretation). Provide no additional explanation after the JSON.’I Ca (HVA)’, ‘I Ca (Rare)’, ‘I Ca (T-type LT)’, ‘I H’, ‘I K (A-type)’, ‘I K (Ca-activated)’, ‘I K (Delayed Rectifier)’, ‘I K (M-type)’, ‘I K (Rare)’, ‘I Na (Persistent)’, ‘I Na (Rare)’, ‘I Na (Slow inactivation)’, ‘I Na (Transient)’, ‘I Other (Rare)’, ‘R GABA’, ‘R Glutamate (AMPA)’, ‘R Glutamate (NMDA)’, ‘R Other (Rare)’, ‘Z Neither’

**Table S1.**
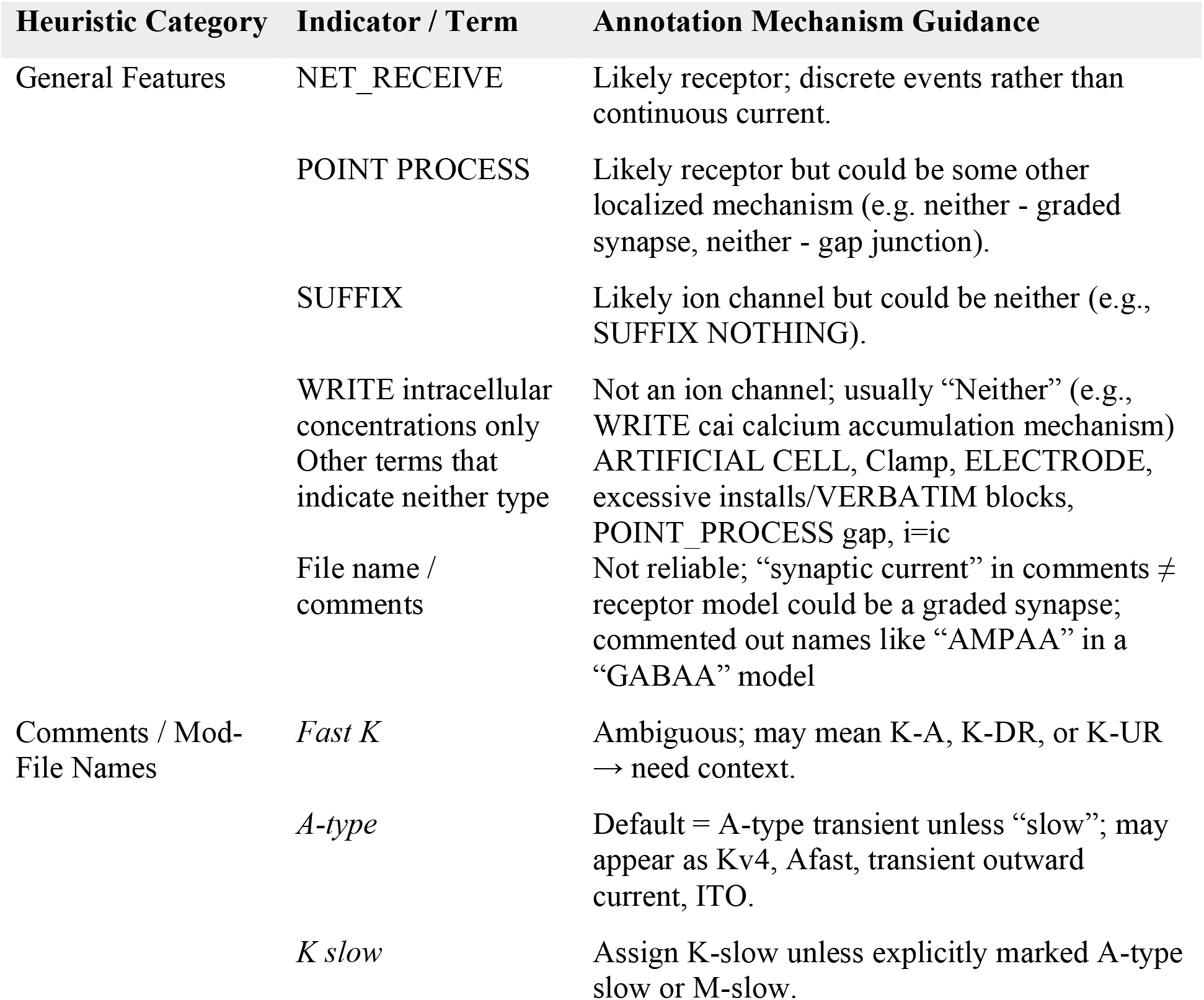
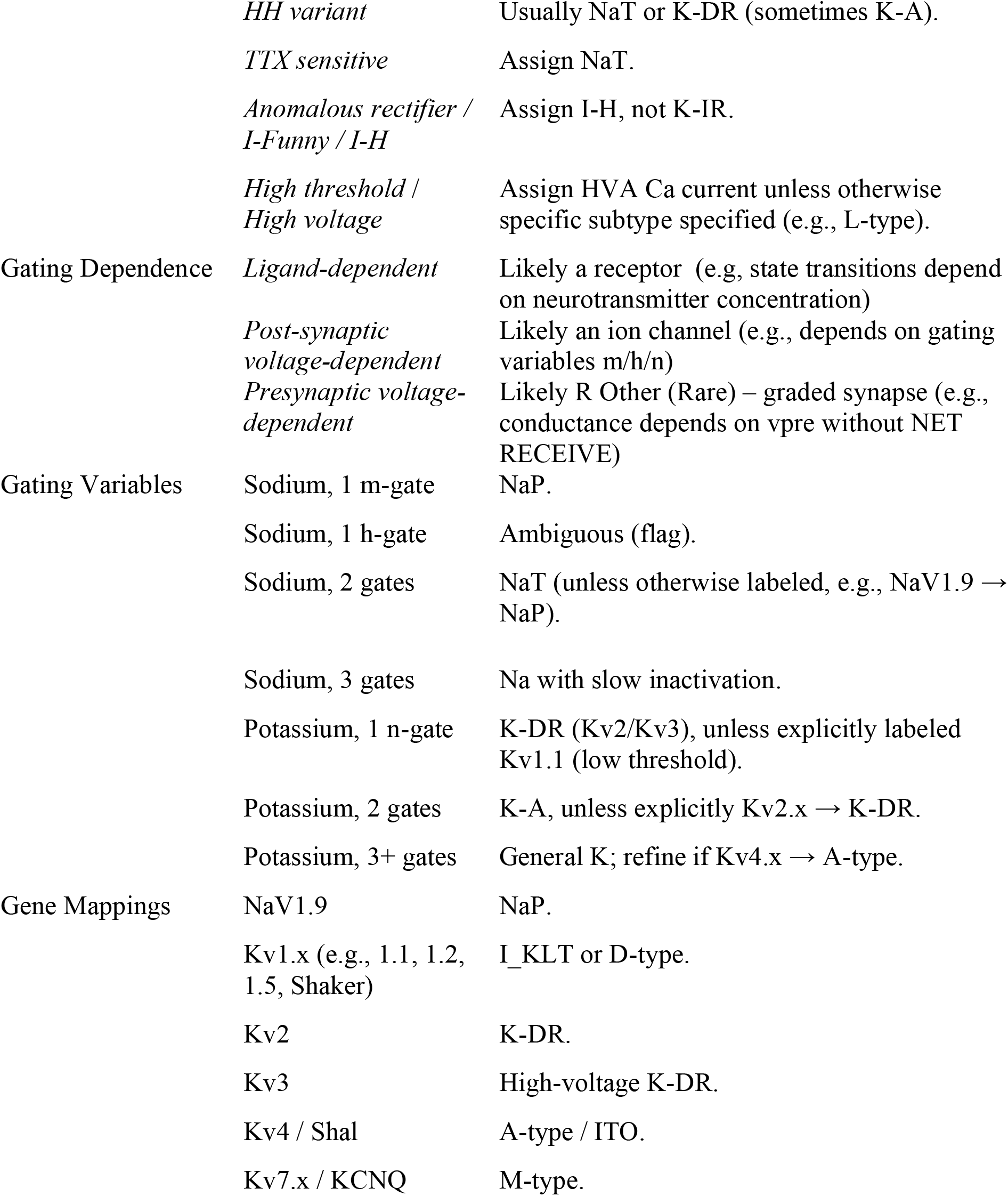
Annotation Heuristics for Manual Labeling of ModelDB Files.

**Table S2.**
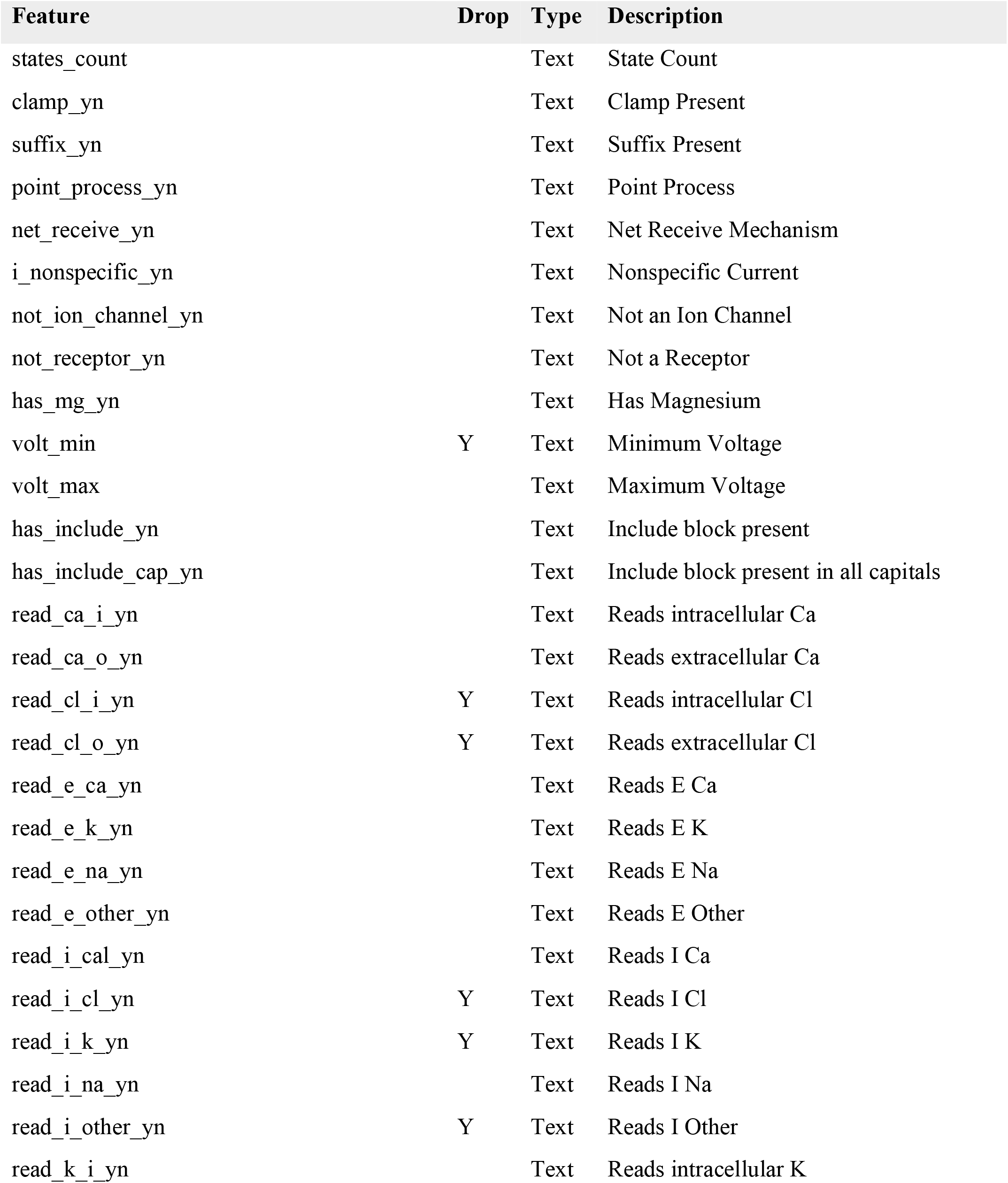
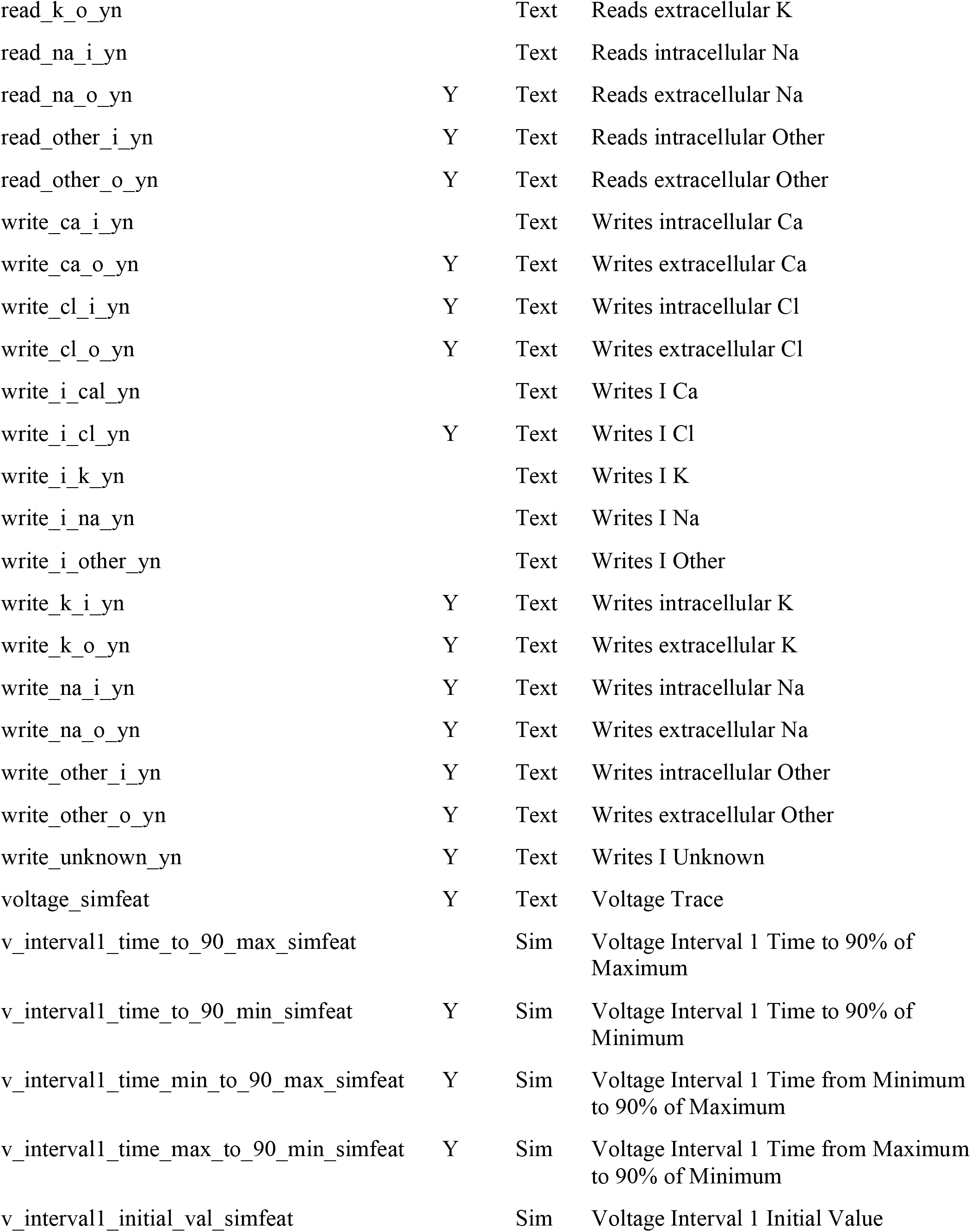
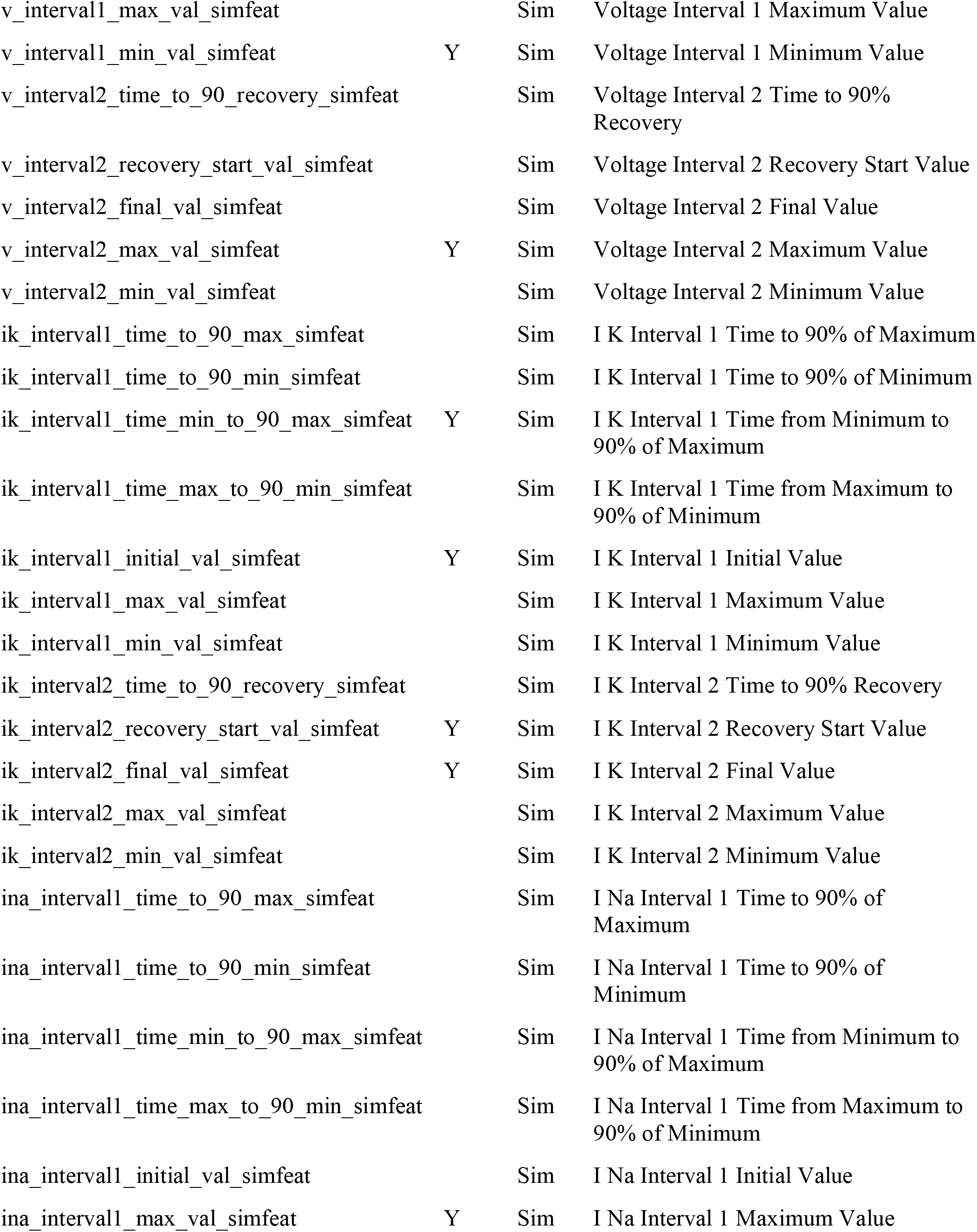
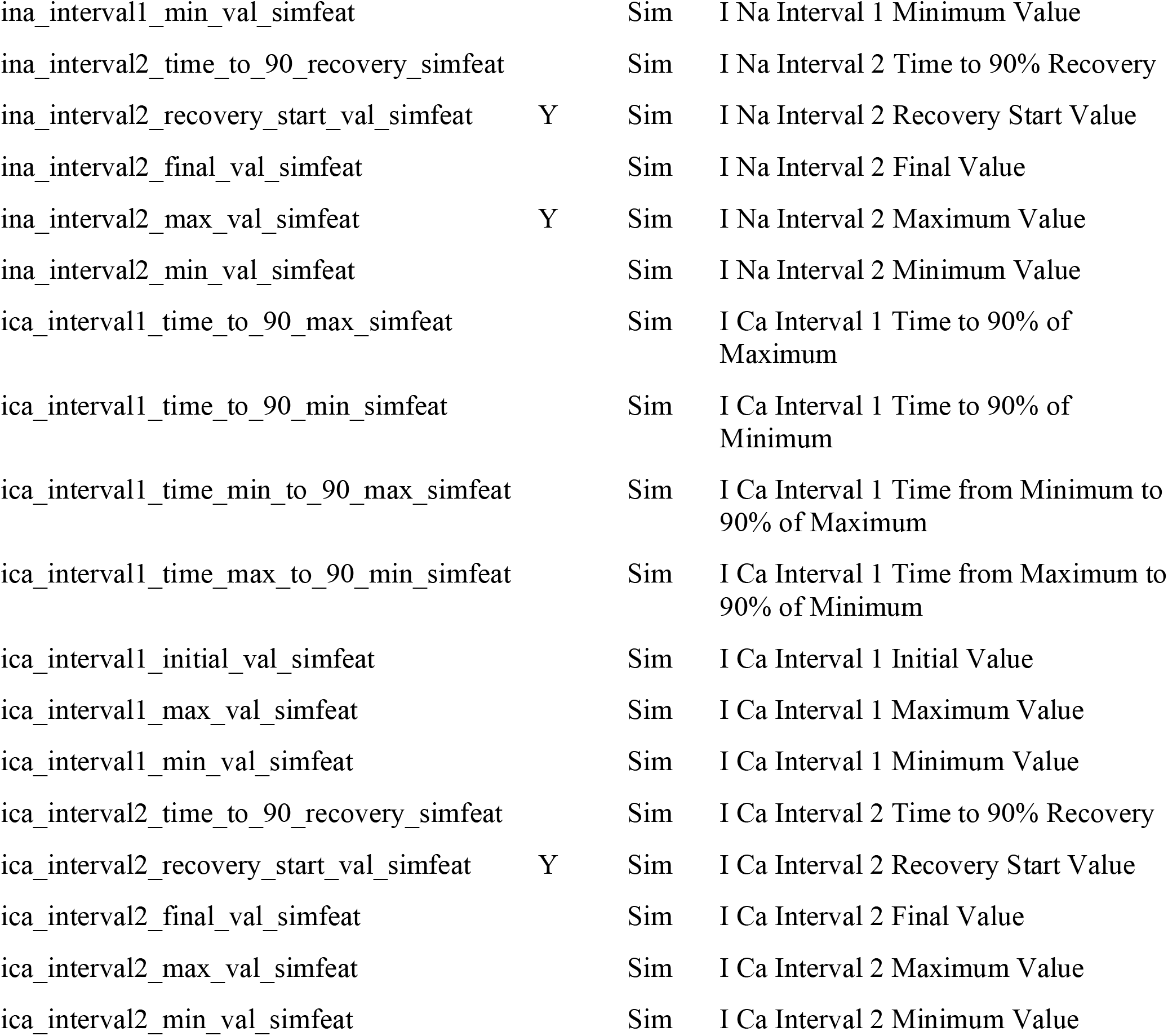
Feature List for XGBoost.

| Feature | Drop | Type | Description |
| --- | --- | --- | --- |
| states_count |  | Text | State Count |
| clamp_yn |  | Text | Clamp Present |
| suffix_yn |  | Text | Suffix Present |
| point_process_yn |  | Text | Point Process |
| net_receive_yn |  | Text | Net Receive Mechanism |
| i_nonspecific_yn |  | Text | Nonspecific Current |
| not_ion_channel_yn |  | Text | Not an Ion Channel |
| not_receptor_yn |  | Text | Not a Receptor |
| has_mg_yn |  | Text | Has Magnesium |
| volt_min | Y | Text | Minimum Voltage |
| volt_max |  | Text | Maximum Voltage |
| has_include_yn |  | Text | Include block present |
| has_include_cap_yn |  | Text | Include block present in all capitals |
| read_ca_i_yn |  | Text | Reads intracellular Ca |
| read_ca_o_yn |  | Text | Reads extracellular Ca |
| read_cl_i_yn | Y | Text | Reads intracellular Cl |
| read_cl_o_yn | Y | Text | Reads extracellular Cl |
| read_e_ca_yn |  | Text | Reads E Ca |
| read_e_k_yn |  | Text | Reads E K |
| read_e_na_yn |  | Text | Reads E Na |
| read_e_other_yn |  | Text | Reads E Other |
| read_i_cal_yn |  | Text | Reads I Ca |
| read_i_cl_yn | Y | Text | Reads I Cl |
| read_i_k_yn | Y | Text | Reads I K |
| read_i_na_yn |  | Text | Reads I Na |
| read_i_other_yn | Y | Text | Reads I Other |
| read_k_i_yn |  | Text | Reads intracellular K |
| read_k_o_yn |  | Text | Reads extracellular K |
| read_na_i_yn |  | Text | Reads intracellular Na |
| read_na_o_yn | Y | Text | Reads extracellular Na |
| read_other_i_yn | Y | Text | Reads intracellular Other |
| read_other_o_yn | Y | Text | Reads extracellular Other |
| write_ca_i_yn |  | Text | Writes intracellular Ca |
| write_ca_o_yn | Y | Text | Writes extracellular Ca |
| write_cl_i_yn | Y | Text | Writes intracellular Cl |
| write_cl_o_yn | Y | Text | Writes extracellular Cl |
| write_i_cal_yn |  | Text | Writes I Ca |
| write_i_cl_yn | Y | Text | Writes I Cl |
| write_i_k_yn |  | Text | Writes I K |
| write_i_na_yn |  | Text | Writes I Na |
| write_i_other_yn |  | Text | Writes I Other |
| write_k_i_yn | Y | Text | Writes intracellular K |
| write_k_o_yn | Y | Text | Writes extracellular K |
| write_na_i_yn | Y | Text | Writes intracellular Na |
| write_na_o_yn | Y | Text | Writes extracellular Na |
| write_other_i_yn | Y | Text | Writes intracellular Other |
| write_other_o_yn | Y | Text | Writes extracellular Other |
| write_unknown_yn | Y | Text | Writes I Unknown |
| voltage_simfeat | Y | Text | Voltage Trace |
| v_intervall_time_to_90_max_simfeat |  | Sim | Voltage Interval 1 Time to 90% of Maximum |
| v_intervall_time_to_90_min_simfeat | Y | Sim | Voltage Interval 1 Time to 90% of Minimum |
| v_intervall_time_min_to_90_max_simfeat | Y | Sim | Voltage Interval 1 Time from Minimum to 90% of Maximum |
| v_intervall_time_max_to_90_min_simfeat | Y | Sim | Voltage Interval 1 Time from Maximum to 90% of Minimum |
| v_intervall_initial_val_simfeat |  | Sim | Voltage Interval 1 Initial Value |
| v_interval1_max_val_simfeat |  | Sim | Voltage Interval 1 Maximum Value |
| v_interval1_min_val_simfeat | Y | Sim | Voltage Interval 1 Minimum Value |
| v_interval2_time_to_90_recovery_simfeat |  | Sim | Voltage Interval 2 Time to 90% Recovery |
| v_interval2_recovery_start_val_simfeat |  | Sim | Voltage Interval 2 Recovery Start Value |
| v_interval2_final_val_simfeat |  | Sim | Voltage Interval 2 Final Value |
| v_interval2_max_val_simfeat | Y | Sim | Voltage Interval 2 Maximum Value |
| v_interval2_min_val_simfeat |  | Sim | Voltage Interval 2 Minimum Value |
| ik_interval1_time_to_90_max_simfeat |  | Sim | I K Interval 1 Time to 90% of Maximum |
| ik_interval1_time_to_90_min_simfeat |  | Sim | I K Interval 1 Time to 90% of Minimum |
| ik_interval1_time_min_to_90_max_simfeat | Y | Sim | I K Interval 1 Time from Minimum to 90% of Maximum |
| ik_interval1_time_max_to_90_min_simfeat |  | Sim | I K Interval 1 Time from Maximum to 90% of Minimum |
| ik_interval1_initial_val_simfeat | Y | Sim | I K Interval 1 Initial Value |
| ik_interval1_max_val_simfeat |  | Sim | I K Interval 1 Maximum Value |
| ik_interval1_min_val_simfeat |  | Sim | I K Interval 1 Minimum Value |
| ik_interval2_time_to_90_recovery_simfeat |  | Sim | I K Interval 2 Time to 90% Recovery |
| ik_interval2_recovery_start_val_simfeat | Y | Sim | I K Interval 2 Recovery Start Value |
| ik_interval2_final_val_simfeat | Y | Sim | I K Interval 2 Final Value |
| ik_interval2_max_val_simfeat |  | Sim | I K Interval 2 Maximum Value |
| ik_interval2_min_val_simfeat |  | Sim | I K Interval 2 Minimum Value |
| ina_interval1_time_to_90_max_simfeat |  | Sim | I Na Interval 1 Time to 90% of Maximum |
| ina_interval1_time_to_90_min_simfeat |  | Sim | I Na Interval 1 Time to 90% of Minimum |
| ina_interval1_time_min_to_90_max_simfeat |  | Sim | I Na Interval 1 Time from Minimum to 90% of Maximum |
| ina_interval1_time_max_to_90_min_simfeat |  | Sim | I Na Interval 1 Time from Maximum to 90% of Minimum |
| ina_interval1_initial_val_simfeat |  | Sim | I Na Interval 1 Initial Value |
| ina_interval1_max_val_simfeat | Y | Sim | I Na Interval 1 Maximum Value |
| ina_interval1_min_val_simfeat |  | Sim | I Na Interval 1 Minimum Value |
| ina_interval2_time_to_90_recovery_simfeat |  | Sim | I Na Interval 2 Time to 90% Recovery |
| ina_interval2_recovery_start_val_simfeat | Y | Sim | I Na Interval 2 Recovery Start Value |
| ina_interval2_final_val_simfeat |  | Sim | I Na Interval 2 Final Value |
| ina_interval2_max_val_simfeat | Y | Sim | I Na Interval 2 Maximum Value |
| ina_interval2_min_val_simfeat |  | Sim | I Na Interval 2 Minimum Value |
| ica_interval1_time_to_90_max_simfeat |  | Sim | I Ca Interval 1 Time to 90% of Maximum |
| ica_interval1_time_to_90_min_simfeat |  | Sim | I Ca Interval 1 Time to 90% of Minimum |
| ica_interval1_time_min_to_90_max_simfeat |  | Sim | I Ca Interval 1 Time from Minimum to 90% of Maximum |
| ica_interval1_time_max_to_90_min_simfeat |  | Sim | I Ca Interval 1 Time from Maximum to 90% of Minimum |
| ica_interval1_initial_val_simfeat |  | Sim | I Ca Interval 1 Initial Value |
| ica_interval1_max_val_simfeat |  | Sim | I Ca Interval 1 Maximum Value |
| ica_interval1_min_val_simfeat |  | Sim | I Ca Interval 1 Minimum Value |
| ica_interval2_time_to_90_recovery_simfeat |  | Sim | I Ca Interval 2 Time to 90% Recovery |
| ica_interval2_recovery_start_val_simfeat | Y | Sim | I Ca Interval 2 Recovery Start Value |
| ica_interval2_final_val_simfeat |  | Sim | I Ca Interval 2 Final Value |
| ica_interval2_max_val_simfeat |  | Sim | I Ca Interval 2 Maximum Value |
| ica_interval2_min_val_simfeat |  | Sim | I Ca Interval 2 Minimum Value |

**Table S3.** Pairwise agreement. Counts and percent agreement between model predictions across the full dataset.

| Model 1 | Model 2 | N | Agree n (%) | Disagree n (%) | Kappa |
| --- | --- | --- | --- | --- | --- |
| GPT-5.2 (run 1) | GPT-5.2 (run 2) | 5133 | 4899 (95.4%) | 234 (4.6%) | 0.95 |
| GPT-5.2 (run 1) | GPT-5.2 + heuristics | 5133 | 4713 (91.8%) | 420 (8.2%) | 0.91 |
| GPT-5.2 (run 2) | GPT-5.2 + heuristics | 5133 | 4695 (91.5%) | 438 (8.5%) | 0.91 |
| GPT-5.2 + heuristics | GPT-mini + heuristics | 5133 | 4618 (90.0%) | 515 (10.0%) | 0.89 |
| GPT-mini | GPT-mini + heuristics | 5132 | 4617 (90.0%) | 515 (10.0%) | 0.89 |
| GPT-5.2 (run 2) | GPT-mini + heuristics | 5133 | 4546 (88.6%) | 587 (11.4%) | 0.87 |
| GPT-5.2 (run 1) | GPT-mini + heuristics | 5133 | 4542 (88.5%) | 591 (11.5%) | 0.87 |
| GPT-5.2 (run 2) | GPT-mini | 5132 | 4539 (88.4%) | 593 (11.6%) | 0.87 |
| GPT-5.2 (run 1) | GPT-mini | 5132 | 4527 (88.2%) | 605 (11.8%) | 0.87 |
| GPT-5.2 + heuristics | GPT-mini | 5132 | 4432 (86.4%) | 700 (13.6%) | 0.85 |

**Table S4.** Run-to-run agreement vs. human annotation confidence. Agreement between repeated runs of each model for high- and lower-confidence cases.

| Model (run-to-run) | High Confidence<br>(n=922) | Low Confidence<br>(n=177) | Overall<br>(n=1099) |
| --- | --- | --- | --- |
| GPT-5.2 (run 1 vs. run 2) | 879 (95.3%) | 167 (94.4%) | 1046 (95.2%) |
| GPT-5.2 vs. GPT-5.2 + heuristics | 849 (92.1%) | 160 (90.4%) | 1009 (91.8%) |
| GPT-mini vs. GPT-mini +<br>heuristics | 839 (91.0%) | 145 (81.9%) | 984 (89.5%) |
| GPT-5.2 vs. GPT-mini | 833 (90.3%) | 141 (79.7%) | 974 (88.6%) |

**Figure S1.**
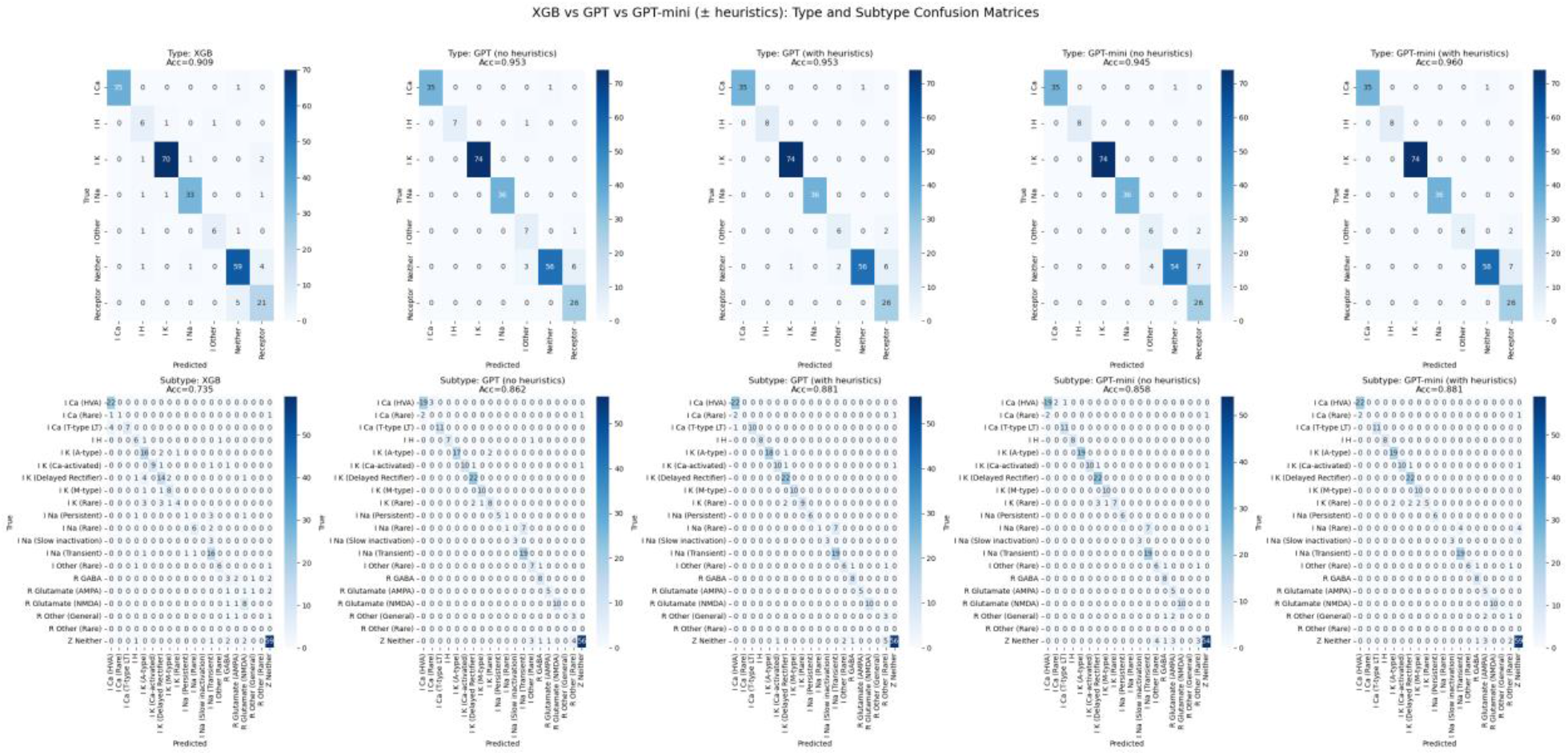
Confusion Matrices. Type and subtype confusion matrices for XGBoost (left), GPT without heuristics (middle), and GPT with heuristics.

**Figure S2.**
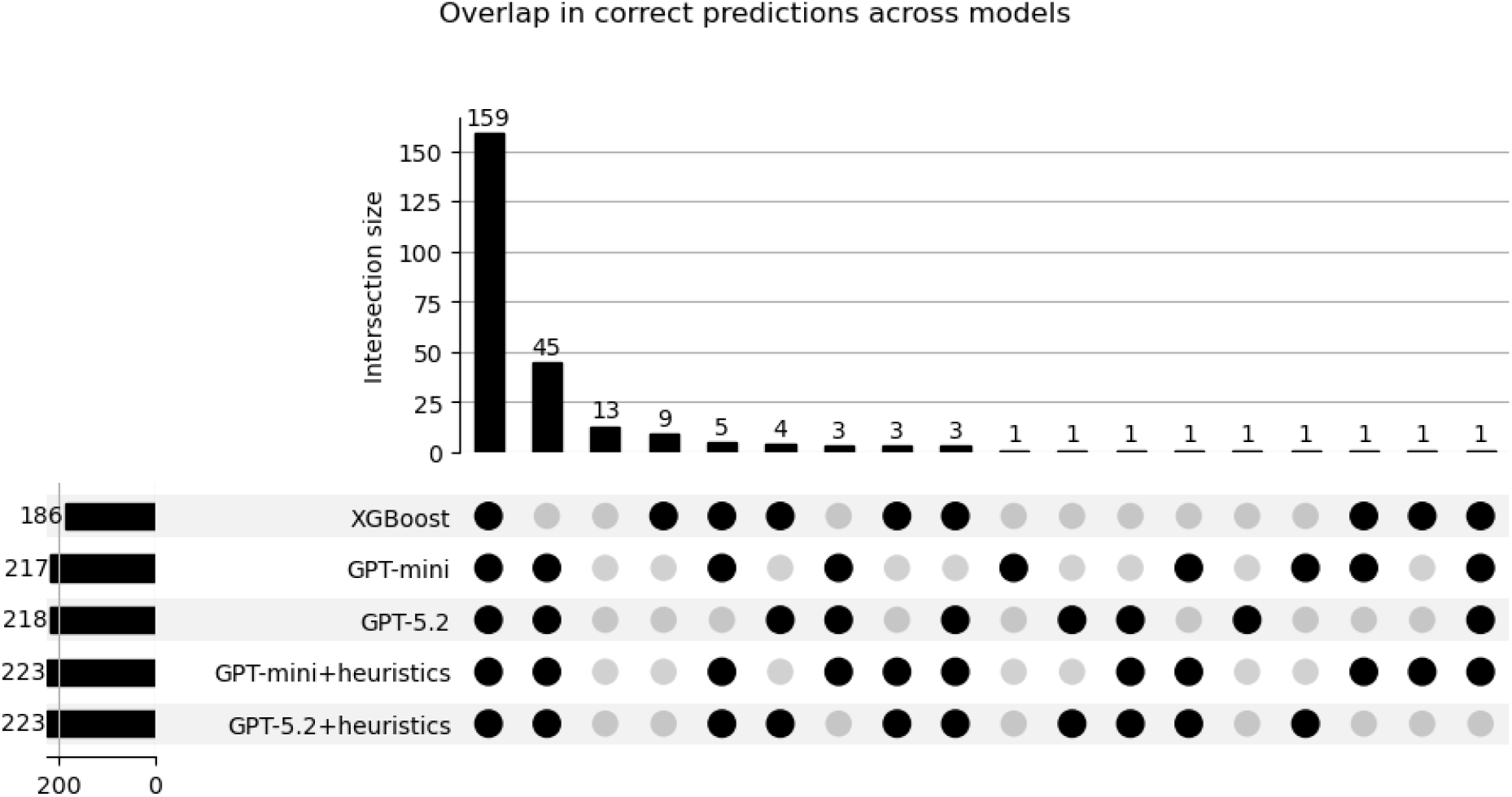
Correct Classification Overlap. UpSet plot showing overlap in correctly classified files across XGBoost and GPT models. Filled dots indicate models included in each intersection, and horizontal bars show the total number of correctly classified files per model. Vertical bars represent the number of files in each intersection.

**Figure S3.**
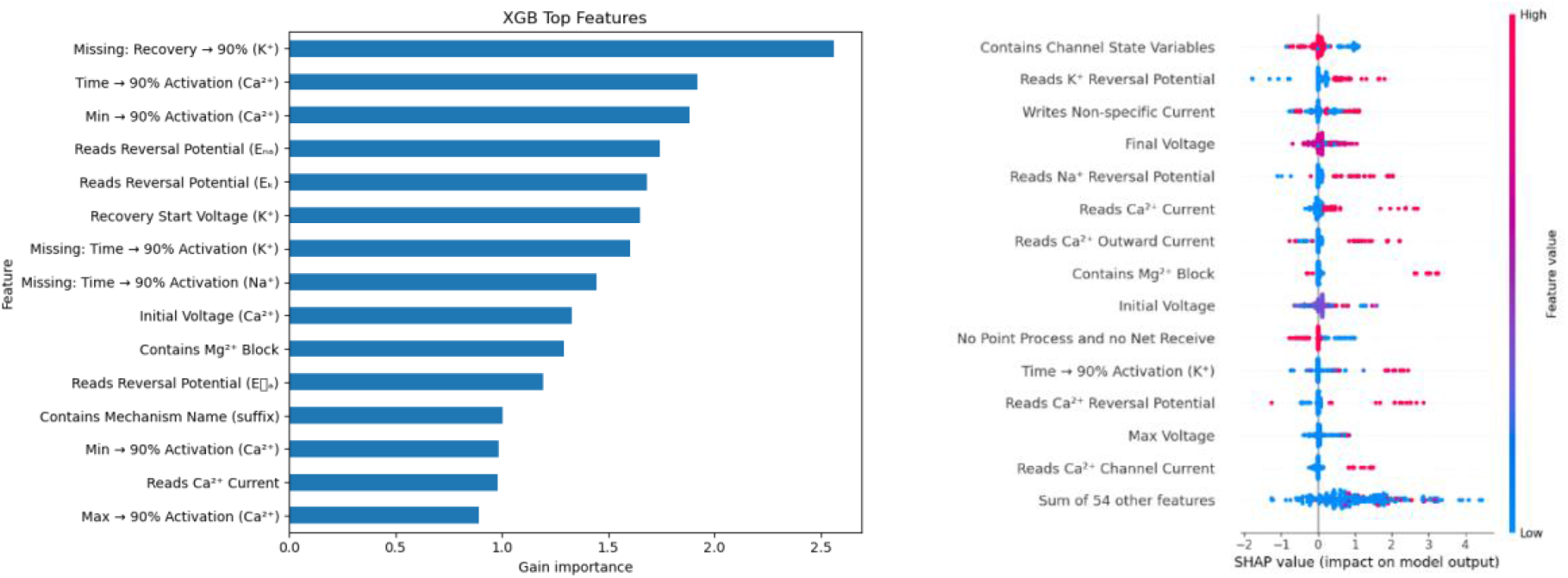
XGBoost Feature Importance. (A) Top 15 features by ranked by XGBoost importance. (B) SHAP summary plot showing the impact of individual features on model output.

